## Supplementary material for "Pupil size reflects activation of subcortical ascending arousal system nuclei during rest": Figure S1, Figure S2, Figure S3, Figure S4

**Supplementary Methods**

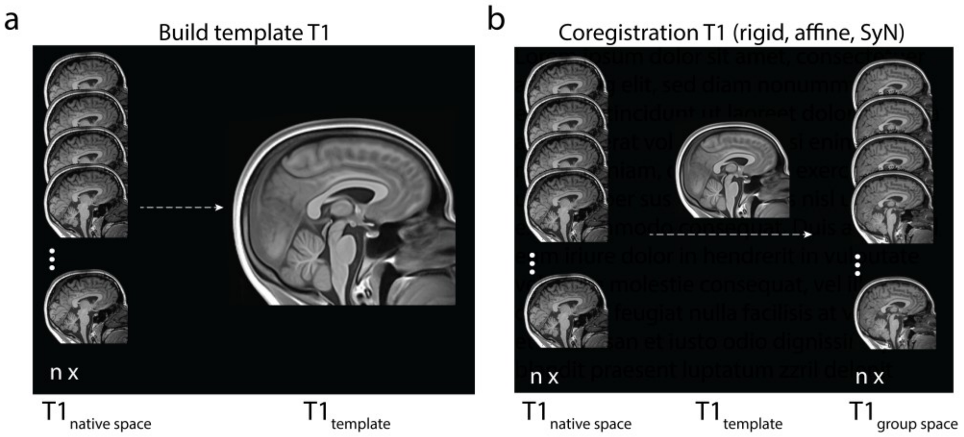


***Figure S1. Overview of template building and coregistration steps performed in ANTs.*** *(a) Individual T1 images ($T1_{native\ space}$) were submitted to a template building algorithm where they were aligned and used to generate a group average ($T1_{template}$). (b) Individual T1 images ($T1_{native\ space}$) were coregistered to the T1 template in a separate coregistration step, generating individual T1 images in group space ($T1_{group\ space}$). Abbreviations: SyN - symmetric normalization.*

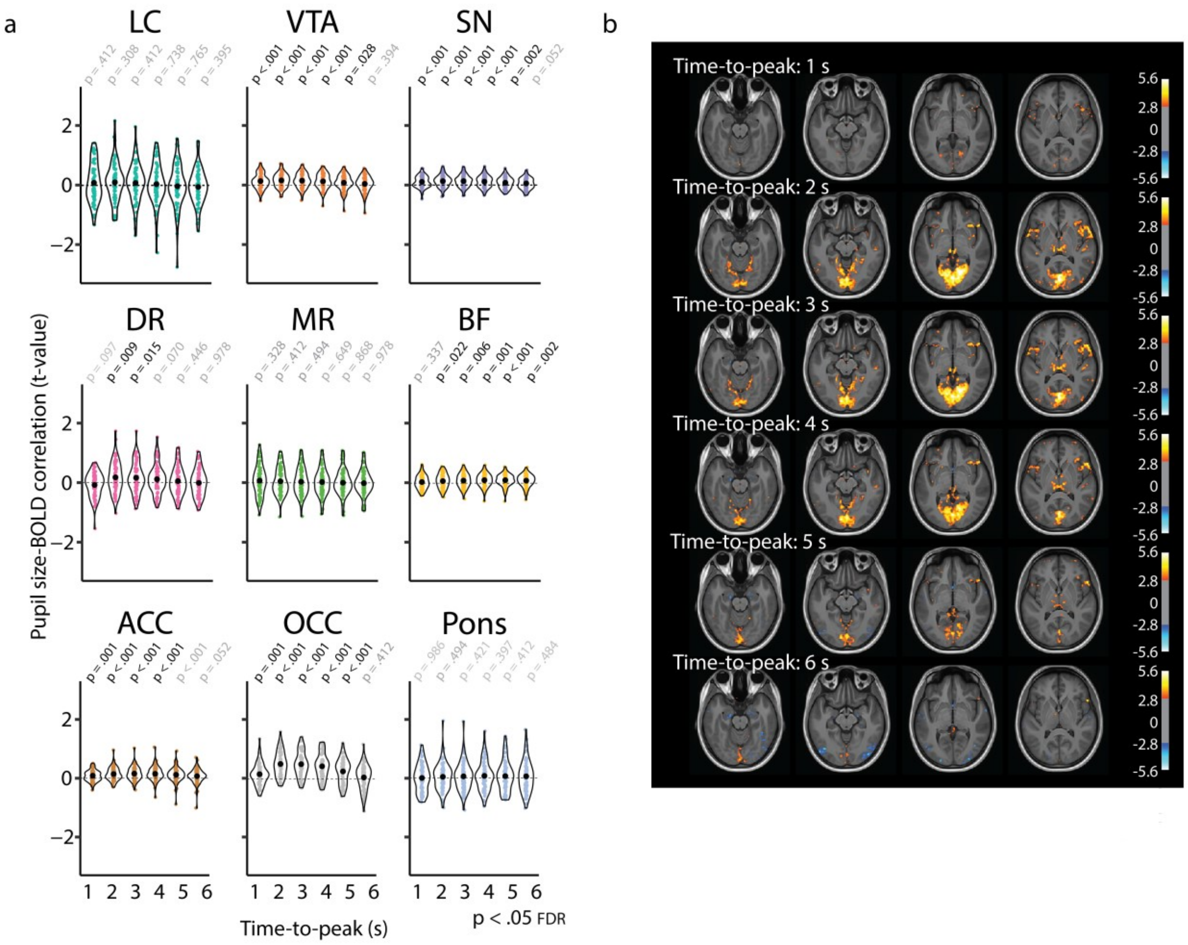


***Figure S2. Pupil derivative-AAS correlations based on systematic adjustment of the time-to-peak.*** *(a) Graphs show the individual t-statistics (black point indicates the mean) of the pupil derivative-BOLD signal correlation as a function of the systematically adjusted TTP for all ROIs, p-values refer to one-sample t-tests (difference from zero; FDR-corrected). (b) The corresponding whole-brain associations between BOLD signal and the pupil derivative as a function of the time-to-peak (TTP) of the HRF (1-s TTP: top panel to 6-s TTP: bottom panel). The warm colours reflect areas in which BOLD activity is positively correlated with pupil derivative. The cold colours reflect areas where BOLD activity is negatively correlated with the pupil derivative. Both positive and negative contrast maps were sampled at p < .005 (uncorrected) for visualization only. Abbreviations: LC – locus coeruleus, VTA – ventral tegmental area, SN – substantia nigra, DR – dorsal raphe, MR – medial raphe, ACC – anterior cingulate cortex, OCC – calcarine sulcus*

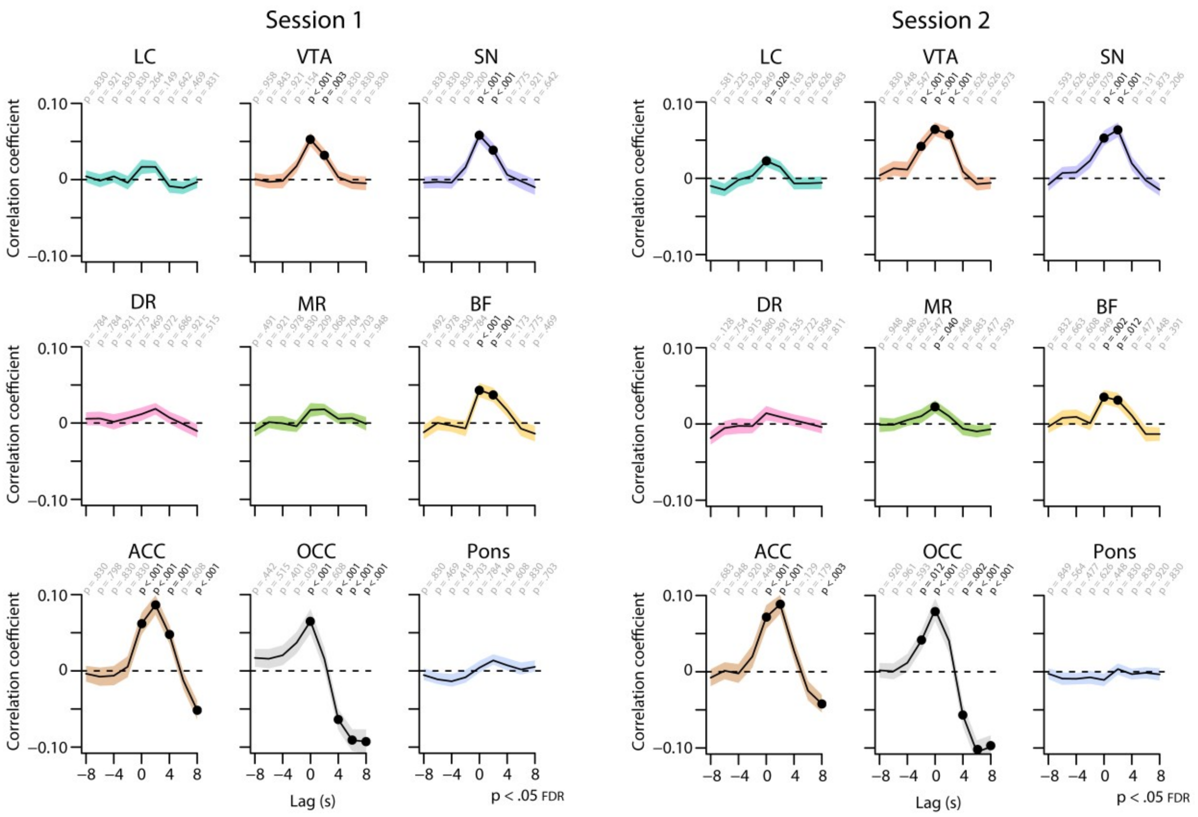


***Figure S3. Cross-correlations between the unconvolved pupil size time series and the BOLD time series at various time lags for each resting-state session.*** *The BOLD time series was shifted forwards and backwards by 8 s in steps of 2 s. Positive (negative) time lags indicate that the pupil signal precedes (follows) the BOLD signal. Black lines indicate the grand mean and shaded regions indicate the standard error of the mean. Black dots indicate significant one-sample t-tests (p < .05, FDR-corrected). Abbreviations: LC – locus coeruleus, VTA – ventral tegmental area, SN – substantia nigra, DR – dorsal raphe, MR – medial raphe, ACC – anterior cingulate cortex, OCC – calcarine sulcus.*

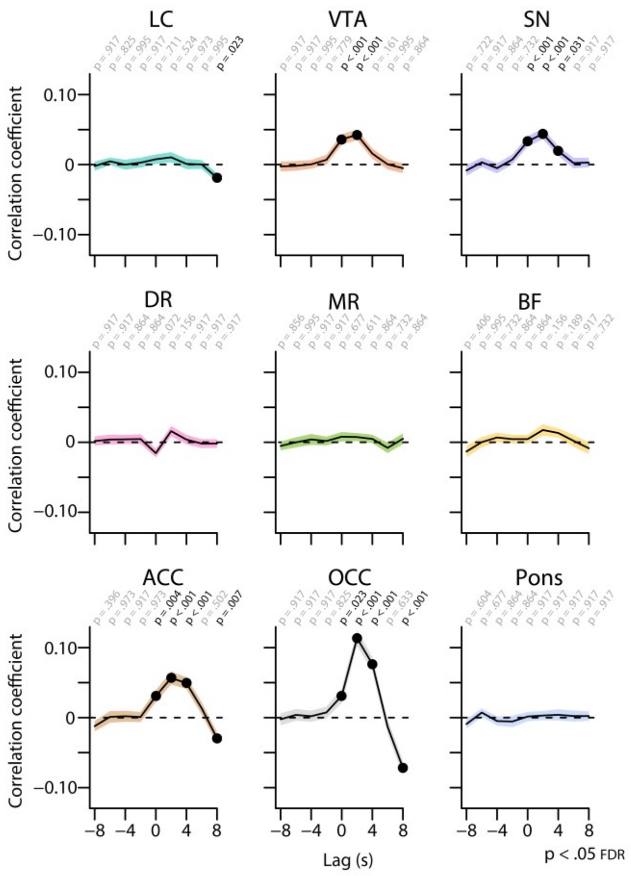


***Figure S4. Cross-correlations between the unconvolved pupil derivative time series and the BOLD time series at various time lags for each resting-state session.*** *The BOLD time series was shifted forwards and backwards by 8 s in steps of 2 s. Positive (negative) time lags indicate that the pupil signal precedes (follows) the BOLD signal. Shaded regions indicate the standard error of the mean. Black dots indicate significant one-sample t-tests (p < .05, FDR-corrected). Abbreviations: LC – locus coeruleus, VTA – ventral tegmental area, SN – substantia nigra, DR – dorsal raphe, MR – medial raphe, ACC – anterior cingulate cortex, OCC – calcarine sulcus.*
